# Systems biology approach identifies functional modules and regulatory hubs related to secondary metabolites accumulation after the transition from autotrophic to heterotrophic growth condition in the microalga

**DOI:** 10.1101/841411

**Authors:** B Panahi, Mohammad Farhadian, Mohammad amin Hejazi

## Abstract

Heterotrophic cultures are among the most promising strategies put forth to overcome the low biomass and secondary metabolites productivity challenge. To shedding light on the underlying molecular mechanisms, weighted gene co-expression network analysis (WGCNA) was integrated with transcriptome meta-analysis, connectivity analysis, functional enrichment, and hub genes identification. Meta-analysis and Functional enrichment analysis demonstrated that with the progress to developmental phases at heterotrophic growth condition, most of the biological processes are up-regulated, which leads to the change of genetic architectures and phenotypic outcomes. WGNCA analysis of meta-genes also created four significant functional modules across logarithmic (LG), transition (TR), and production peak (PR) phases. The expression pattern and connectivity characteristics of the brown module as a non-preserved module vary across LG, TR, and PR phases. Functionally analysis identified “Carotenoid biosynthesis”, “Fatty acid metabolism”, and “Methane metabolism” as enriched pathways in the non-preserved module. Moreover, the integrated approach was applied here, identified some hub genes, such as serine hydroxymethyltransferase (SHMT1) for development of metabolites accumulating strains in microalgae. Our study provided a new insight into underlying mechanisms of metabolite accumulation and opens new avenue for the future applied studies in the microalgae field.

---

**Table 1.** Details of conservation analysis of defined modules at different developmental phases with permutation=200.

| Module name | Module size | medianRank | Zsummary |
| --- | --- | --- | --- |
| Blue | 469 | 2 | 32.56 |
| Brown | 184 | 8 | 4.26 |
| Turquoise | 681 | 3 | 34.39 |
| Yellow | 155 | 1 | 19.05 |

**Table 2.**
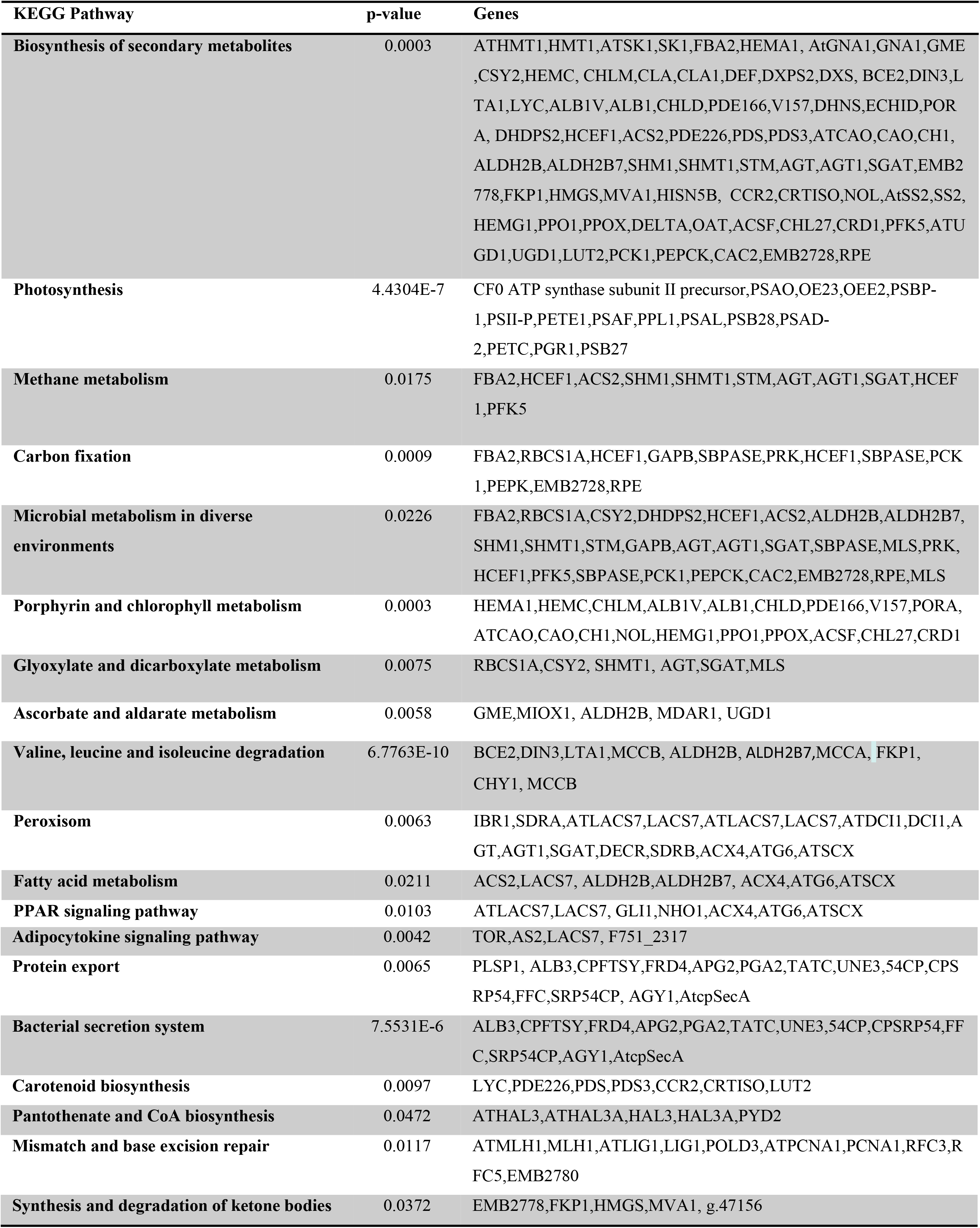
Functional enrichment of non-preserved module based on KEGG database

**Figure 1.**
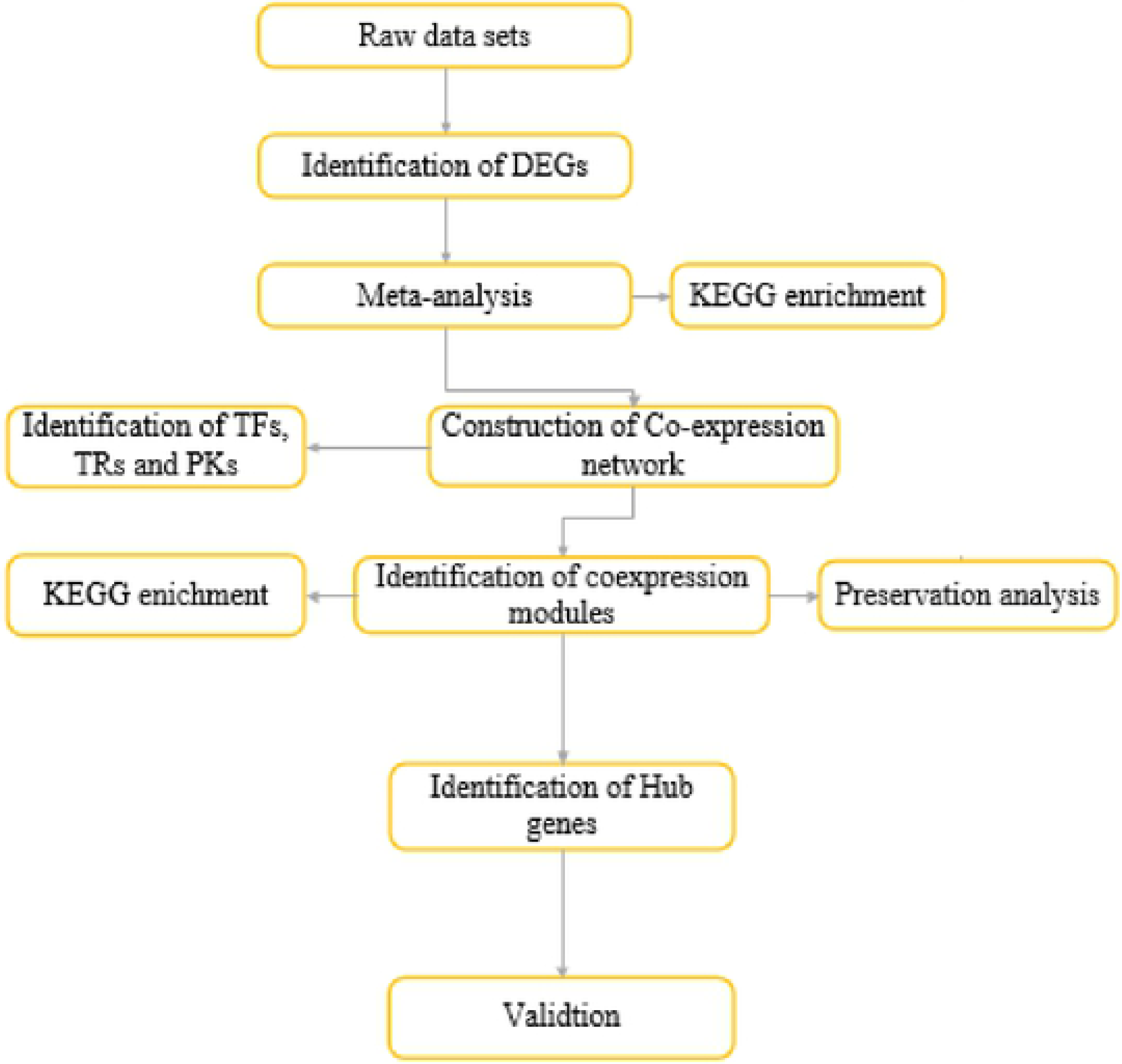
Flow chart of applied systems biology approach in current study.

**Figure 2.**
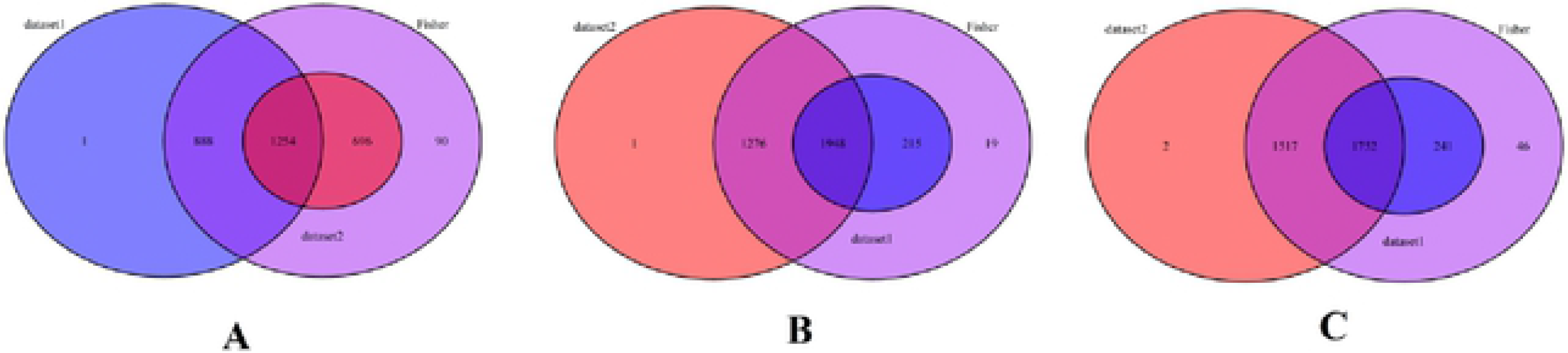
Identified meta genes in two data sets at logarithmic (A), transition (B) and production peak (C) phases after the transition from autotrophic to heterotrophic growth mode.

**Figure 3.**
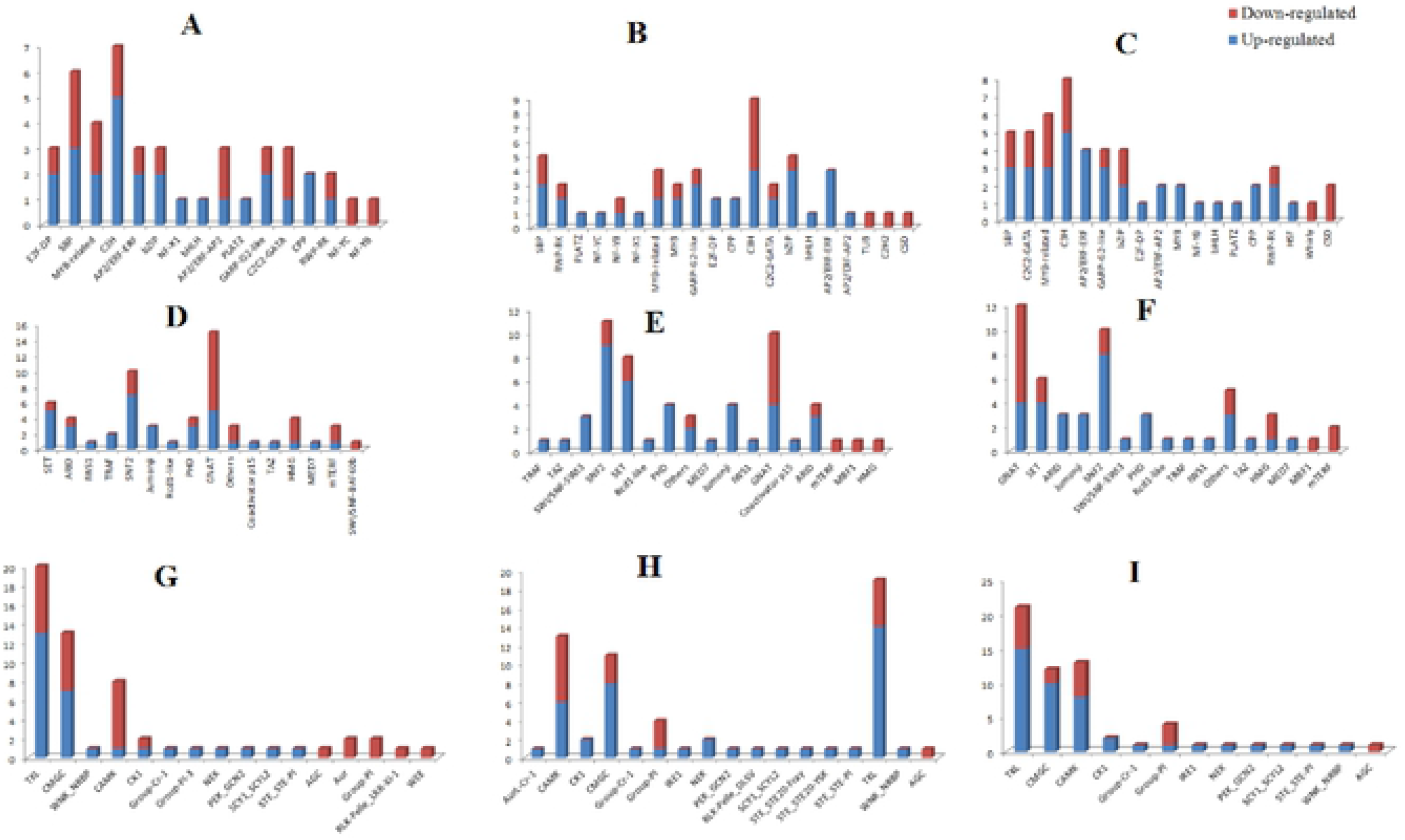
Distribution of TFs, TRs, and PKs families identified in meta genes The number TF families (A, B and C), TRs families (D, E, and F), PKs families (G, H, and I) in meta genes at LG, TR and PR phases, respectively.

**Figure 4.**
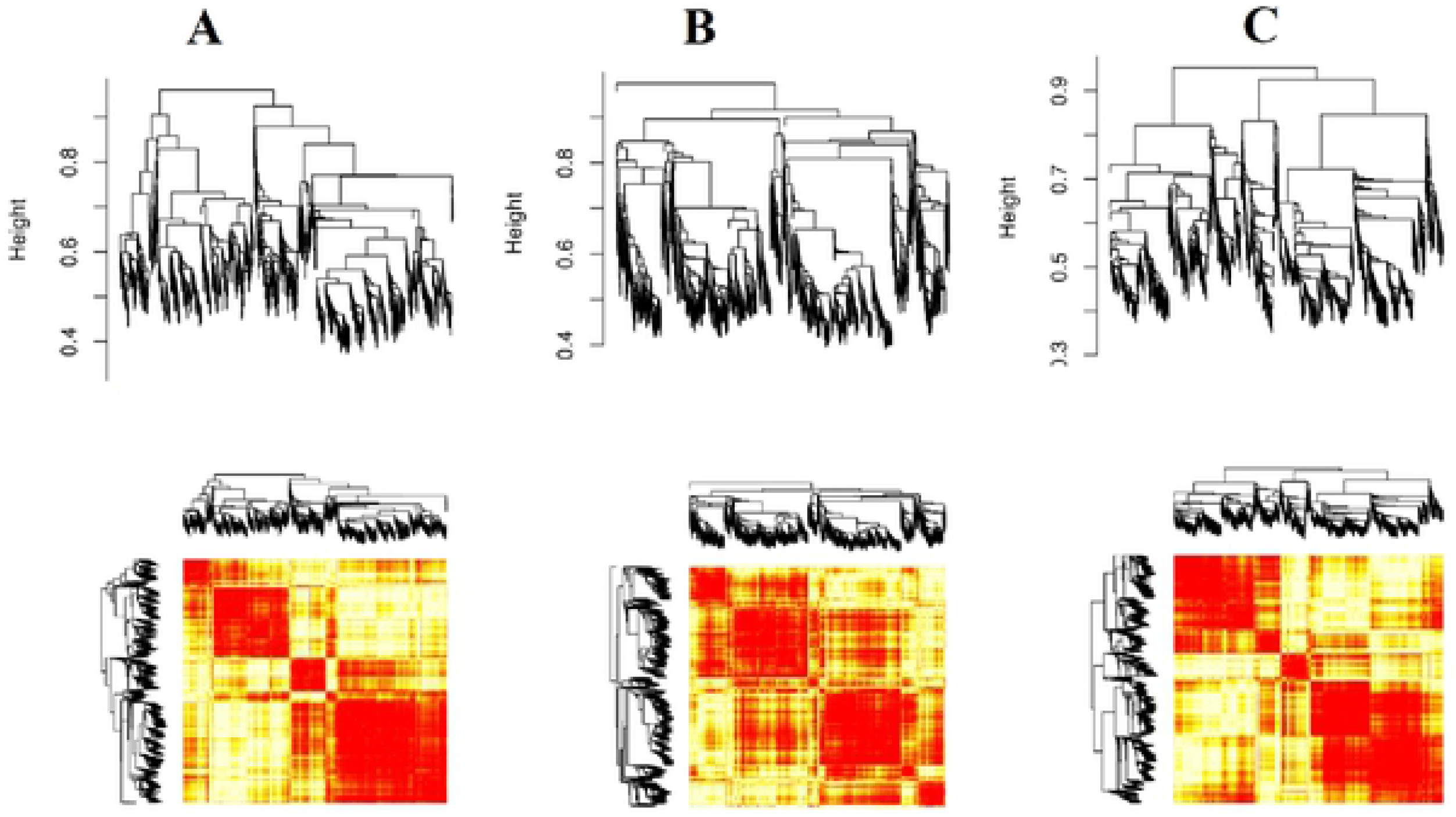
Hierarchical cluster constructed with WGCNA at LG (A), TR (B), and PR (C) phases. Each vertical line (leaf) represents the corresponding genes.

**Figure 5.**
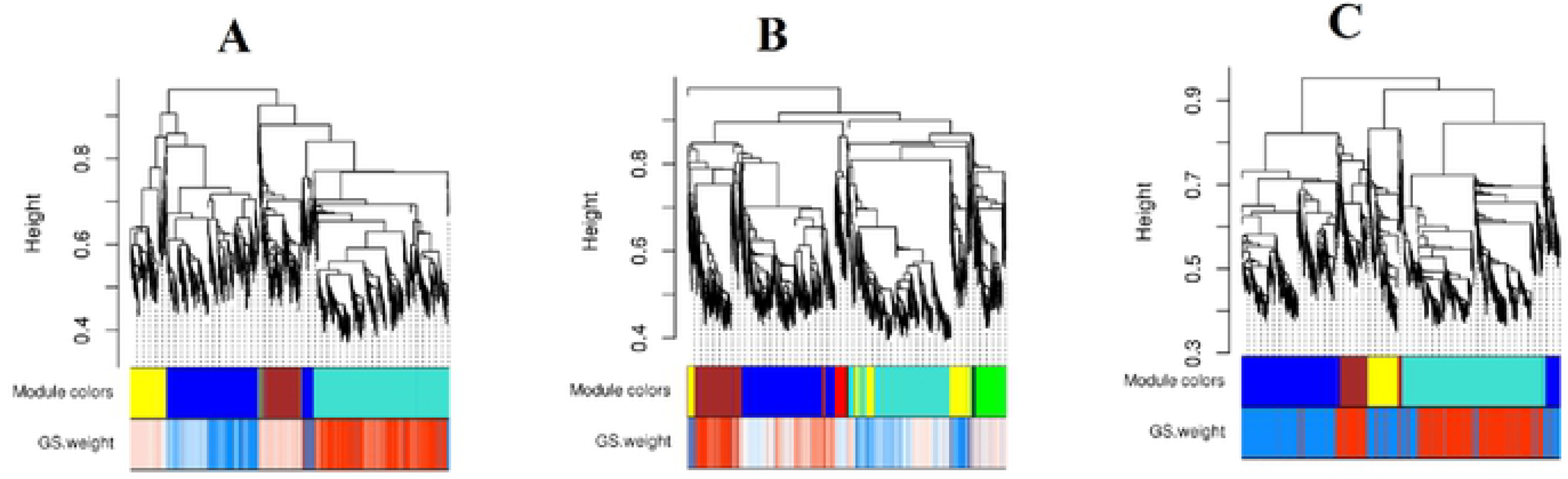
Visual representation of the changes in the module structure between LG (A), TR (B), and PR (C) phases. Modules are illustrated with different colors.

**Figure 6.**
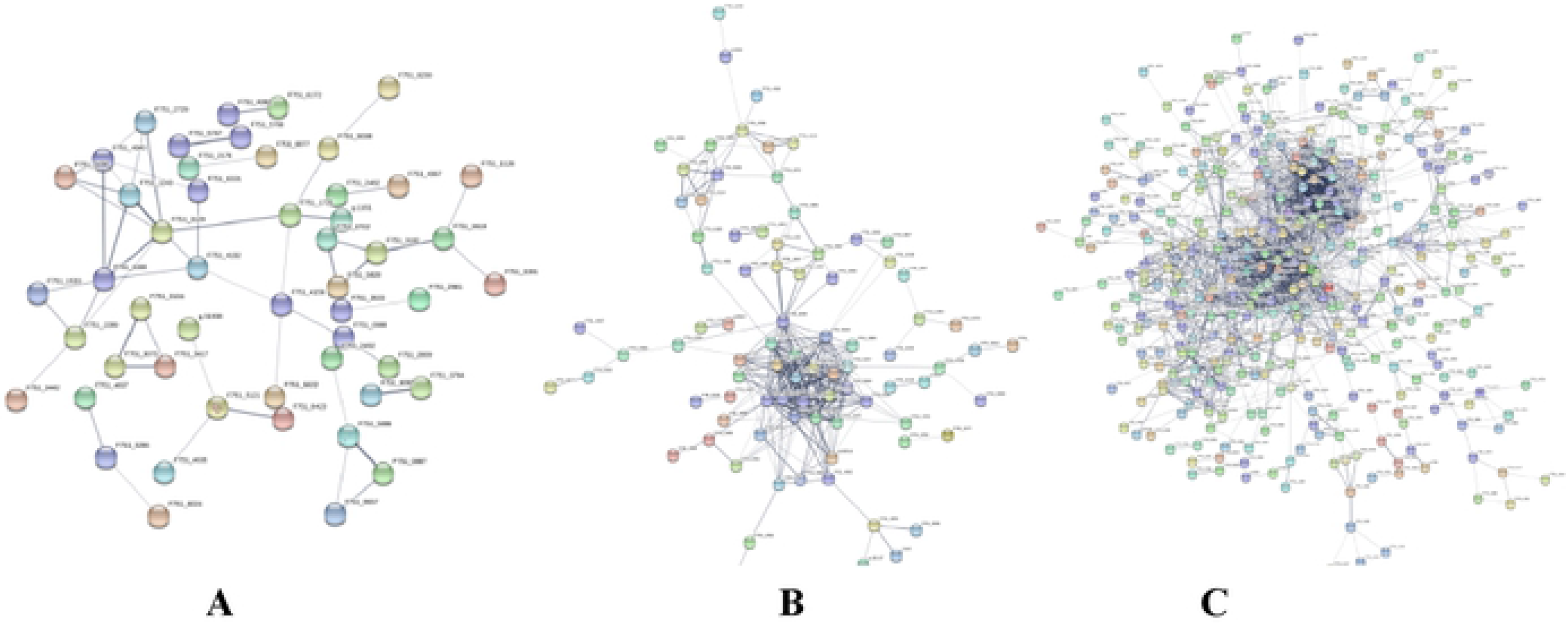
PPI networks of co-expressed meta genes in brown module at LG (A), TR (B), and PR (9C) phases. Changes in intra-module connectivity are highlighted at different phases.

